# Impact of tissue staining and scanner variation on the performance of pathology foundation models: a study of sarcomas and their mimics

**DOI:** 10.1101/2025.08.18.670932

**Authors:** Binghao Chai, Jianan Chen, Paul Cool, Fatine Oumlil, Anna Tollitt, David F. Steiner, Tapabrata Chakraborti, Adrienne M. Flanagan

## Abstract

Histopathological analysis is considered the gold standard for the diagnosis and prognostication of cancer. Recent advances in AI, driven by large-scale digitisation and pan-cancer foundation models, are opening new opportunities for clinical integration. However, it remains unclear how robust these foundation models are to real-world sources of variability, particularly in H&E staining and scanning protocols. In this study, we use soft tissue tumours, a rare and morphologically diverse tumour type, as a challenging test case to systematically investigate the colour-related robustness and generalisability of seven AI models. Controlled staining and scanning experiments were utilised to assess model performance across diverse real-world data sources. Foundation models, particularly UNI-v2, Virchow and TITAN, demonstrated encouraging robustness to staining and scanning variation, particularly when a small number of stain-varied slides were included in the training loop, highlighting their potential as adaptable and data-efficient tools for real-world digital pathology workflows.

## 1 Introduction

Digital pathology refers to the digitisation of whole slide images (WSIs) of stained tissue sections that can be used for diagnostic and research purposes. In recent years, there has been substantial development in the field, driven by advances in scanning technology, computational infrastructure, artificial intelligence (AI) and clinical need [1]. Moving from traditional light microscopy to digital workflows has enabled more scalable and reproducible image analysis, supporting remote consultations and real-time diagnostic decision-making and the use of AI to support pathologists in providing their clinical service. This transition has been accommodated by the widespread adoption of high-speed, high-resolution slide scanners, and improvements in image compression, cloud storage, and digital slide viewers [2]. Traditional AI models have been largely developed for narrowly defined problems, such as mitosis detection [3], nuclei segmentation [4], and Gleason grading in prostate cancer [5], which typically require large amounts of annotated data. In contrast, recent advances in deep learning, particularly the emergence of pathology foundation models, have expanded the scope and flexibility of AI applications [6, 7]. These large-scale models, trained on up to millions of unlabeled image tiles across various tissue types, magnifications, and institutions, are capable of learning multiple general-purpose representations that can be fine-tuned for a variety of downstream tasks [6, 7].

Foundation models can reduce reliance on extensive task-specific annotations, thereby accelerating the introduction of AI into clinical pathology. Such models include Google Path Foundation [8], Virchow [9, 10], and UNI [11] which are general-purpose patch-level encoders, while CONCH is a multimodal foundation model that integrates both image and text data [12]. Slide-level foundation models such as TITAN [13] and PRISM [14] combine patch encoders with pretrained aggregation modules to generate slide-level representations directly. Nevertheless, irrespective of the AI model employed, a consistent high level of performance across WSIs stained using different protocols and scanned on different devices is essential if they are to be introduced into clinical practice, as these two variables can substantially alter image appearance and model behaviour [15–19].

To address the challenges posed by stain variation, researchers have investigated a variety of strategies, including stain normalisation and more generalisable training designs. Stain normalisation (e.g. Reinhard [20], Macenko [21], and Vahadane [22]) aims to standardise colour distributions across slides prior to model training. However, these methods often underperform, particularly in preserving tissue morphology, compared to stain augmentation or deep learning-based normalisation approaches [23– 25]. Deep learning-based stain variation correction methods, for example, those using generative adversarial networks (GANs), achieve better morphological preservation [26–31]. However, these tools typically require large-scale training datasets and tend to generalise poorly to slides from unseen sources. Foundation models have recently emerged as a promising solution to staining and scanning variability. Pretrained on large and diverse datasets, they offer a more robust and generalisable starting point for downstream task adaptation [8, 11–13]. In general, there has been a shift away from extensive stain preprocessing. Data are now often used in their original unnormalised form, due to the computational cost and practical challenges of applying stain normalisation on WSIs [23, 32]. However, the robustness of foundation models to stain and scanner variation remains relatively underexplored.

The aim of this study was to evaluate the extent to which pre-trained foundation models remain robust to staining and scanning variation across real-world images, thereby informing their utility in generalisable digital pathology workflows. We have employed sarcomas and their mimics, a rare and morphologically diverse tumour group [33], to investigate these questions.

## 2 Results

### 2.1 Evaluation of foundation models for slide-level subtyping

All six foundation models (Google Path Foundation [8], UNI-v2 [11], CONCHv1.5 [12], TITAN [13], Virchow [9] and PRISM [14]) along with a ResNet50-based convolutional neural network (CNN) [34] demonstrated a high level of diagnostic accuracy (i.e., the proportion of correctly predicted diagnoses) when trained and tested on an image collection stained in one laboratory and scanned on one device, Cohort 1 (Fig. 1 **a**). Foundation models, TITAN, UNI-v2 and Virchow achieved the highest accuracy, and all models outperformed ResNet50 (Fig. 2 **b**). The performance of all foundation models dropped when tested on Cohort 4-Test, a ‘real-world’ dataset stained in multiple institutions and scanned on a single platform (Fig. 2 **b**). This diagnostic performance across 14 sarcomas and their mimics (Fig. 1 **b**) assessed on the seven models was reflected in the classwise accuracy, with Cohort 4-Test showing a wider variance compared to Cohort 1 (Fig. 1 **c**). The radar plots of the 14 tumour types highlight the difference in performance between Cohorts 1 and 4. In Cohort 4, accuracy was retained for dermatofibrosarcoma protuberans (DFSP), desmoid fibromatosis, and synovial sarcoma, but was reduced moderately for solitary fibrous tumour. Nodular fasciitis and intramuscular myxoma were the lowest performing diagnoses. ResNet50CNN underperformed across all models when tested on both Cohorts 1 and 4 (Fig. 1 **d, e**).

**Fig. 1.**
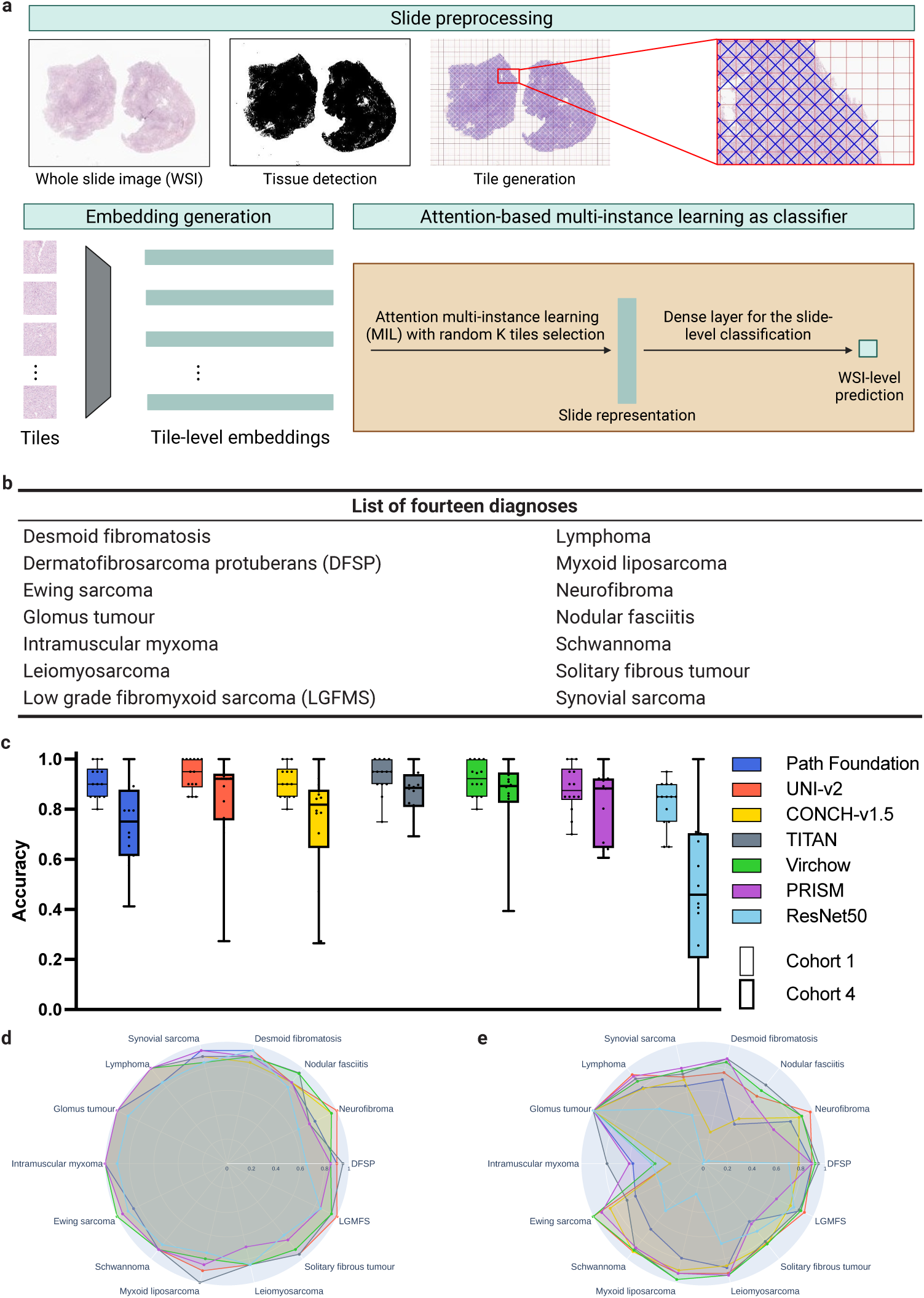
The computational pipeline and subtyping performance for soft tissue tumours. a, WSIs were preprocessed with tissue detection and tile generation on hematoxylin and eosin (H&E)-stained slides using TRIDENT [35, 36]. Tile-level embeddings were generated using six foundation models and ResNet50-CNN before being sent to the attention-based multi-instance learning (MIL) classifier for training. b, The list of 14 diagnoses containing sarcomas and their mimics in this study. c, Class-wise accuracy distribution for Cohort 1-Test and Cohort 4-Test sets across all seven models: Each pair of boxes represents the diagnostic accuracy of the 14 different diagnoses per model. Google Path Foundation (Cohort 1: 0.911 ± 0.066, Cohort 4: 0.736 ± 0.157), UNI-v2 (Cohort 1: 0.939 ± 0.063, Cohort 4: 0.842 ± 0.189), CONCH-v1.5 (Cohort 1: 0.911 ± 0.066, Cohort 4: 0.737 ± 0.233), TITAN (Cohort 1: 0.932 ± 0.070, Cohort 4: 0.869 ± 0.086), Virchow (Cohort 1: 0.921 ± 0.067, Cohort 4: 0.859 ± 0.152), PRISM (Cohort 1: 0.882 ± 0.093, Cohort 4: 0.811 ± 0.140) and ResNet50-CNN (Cohort 1: 0.821 ± 0.091, Cohort 4: 0.466 ± 0.306). Each box represents the 14 subtype accuracies per model. d, e, Radar plots showing the diagnosis-level performance for all seven models on Cohort 1-Test and Cohort 4-Test, respectively. An interactive version of the radar plots is provided (Data Availability section).

**Fig. 2.**
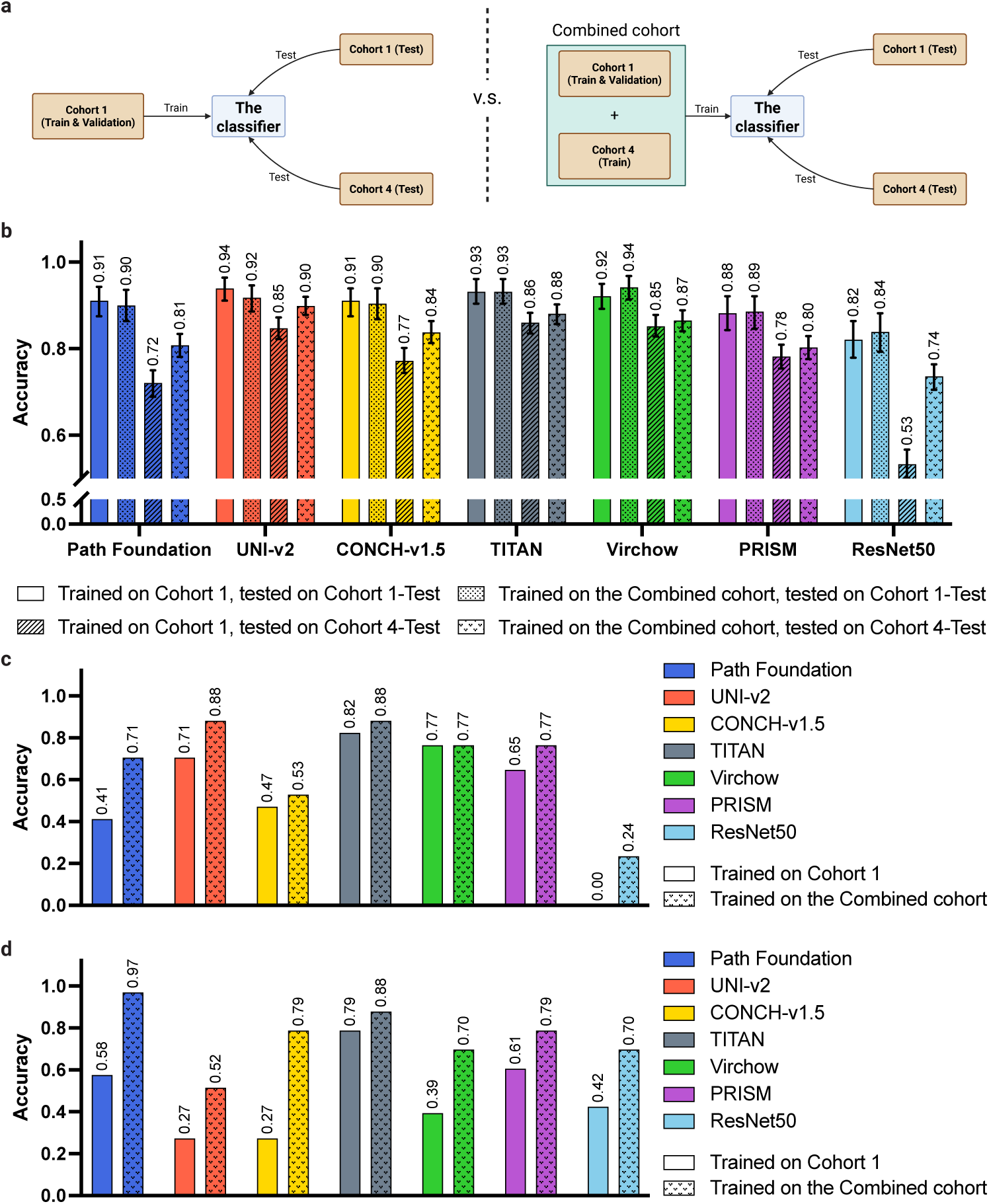
Improvement of classification of 14 soft tissue tumour types by training on WSIs stained in more than one laboratory. a, Experimental design to explore the impact of staining variation under two settings. In the original setting (left), the classifier was trained only on the internal cohort (Cohort 1-Train), and then on a Combined cohort (right) containing an additional 121 WSIs from the multi-institutional cohort (Cohort 4-Train). Both versions of the classifier were evaluated independently on the internal (Cohort 1-Test) and the ’real world’ (Cohort 4-Test) hold-out set. b, The analysis was conducted on Path Foundation, UNI-v2, CONCH-v1.5, TITAN, Virchow, PRISM and ResNet50-CNN models. Models trained only on Cohort 1 revealed a statistically significant decline in classification performance when models were applied to the ’real world’ test cohort (Cohort 4-Test) compared to the internal cohort (Cohort 1-Test): Google Path Foundation (p < 0.001), UNI-v2 (p < 0.001), CONCH-v1.5 (p < 0.001), TITAN (p = 0.004), Virchow (p = 0.006), PRISM (p < 0.001), and ResNet50 (p < 0.001) with chi-squared test. However, when the models were trained on the Combined cohort, a statistically significant improvement in Cohort 4-Test classification performance was observed for Google Path Foundation (p < 0.001), UNI-v2 (p < 0.002), CONCH-v1.5 (p < 0.001), and ResNet50 (p < 0.001); while Virchow (p = 0.478) and slide encoders (TITAN: p = 0.238, PRISM: p = 0.329), not significant. c, d, Performance comparison between ’trained on Cohort 1’ and trained on the Combined cohort’ on Cohort 4-Test set for those failed subtypes (nodular fasciitis (c) and intramuscular myxoma (d), respectively) in the original training setting.

To address the limitations of AI classification models when WSIs are stained in a single institution, we added a subset of externally stained WSIs from Cohort 4Train (n=121, Table 1) to the Cohort 1-Train (Combined cohort, Fig. 2 **a**). This resulted in all patch-based foundation models exhibiting substantial overall accuracy improvements when tested on Cohort 4-Test (Fig. 2 **b**, Table B1). This also led to improved classwise accuracy, and it is noteworthy that results for nodular fasciitis and intramuscular myxoma were better than in the previous experiments (Fig. 2 **c, d**). Interestingly, a minor decline in the performance of Google Path Foundation, UNIv2, and CONCH-v1.5 was observed when the model was tested on Cohort-1 Test but trained on the Combined cohort (Fig. 2 **b**). Full results are reported in Extended Table B3.

**Table 1.** Four collections of WSI employed in this study.

| Dataset | Slides-Total | Slides-Train | Slides-Val | Slides-Test | Stain Institute | Scan Device |
| --- | --- | --- | --- | --- | --- | --- |
| Cohort 1 | 1,394 | 835 | 279 | 280 | Lab A | Aperio |
| Cohort 2 | 1,243 | - | - | 1,243 | Multiple | Multiple |
| Cohort 3 | 335 | - | - | 335 | Lab A | 3 |
| Cohort 4 | 934 | 121 | - | 813 | Multiple | Aperio |
| **Total** | **3,906** | **956** | **279** | **2,671** | **-** | **-** |

### 2.2 Impact of staining and scanning variation

We next attempted to disentangle how differences in staining and scanning contribute to colour variation (Fig. 3; Cohort 2, Table 0). Fig. 4 **a** illustrates the staining and scanning variations using a case of synovial sarcoma. We extracted stain vectors from randomly sampled 3000 *×* 3000 pixel tiles using the Vahadane method [22], which separates the optical density matrix into stain vectors and stain concentration maps. We then performed principal component analysis (PCA) to project the colour vectors in a two-dimensional space (Fig. 4 **b**). Notably, the tile representations on PCA clustered largely by scanning device rather than staining institution. The Aperio-scanned slides (circles) formed a tight cluster in the top right region of the plot, indicating relatively consistent colour vector characteristics, whereas 3D-Histech-scanned slides (triangles) were widely dispersed, reflecting higher intra-scanner colour variability. Hamamatsu-scanned slides (squares, top left region) showed clustering intermediate between the other two scanners (see Fig. C3 for PCA plot with each scanner). These findings support our previous results which showed that variation in staining reduces performance when classifying tumours using AI models, and that the addition of different scanning devices compounds the challenge. The wide distribution across the PCA does not allow quantitative assessment of the impact of staining and scanning of colour variation.

**Fig. 3.**
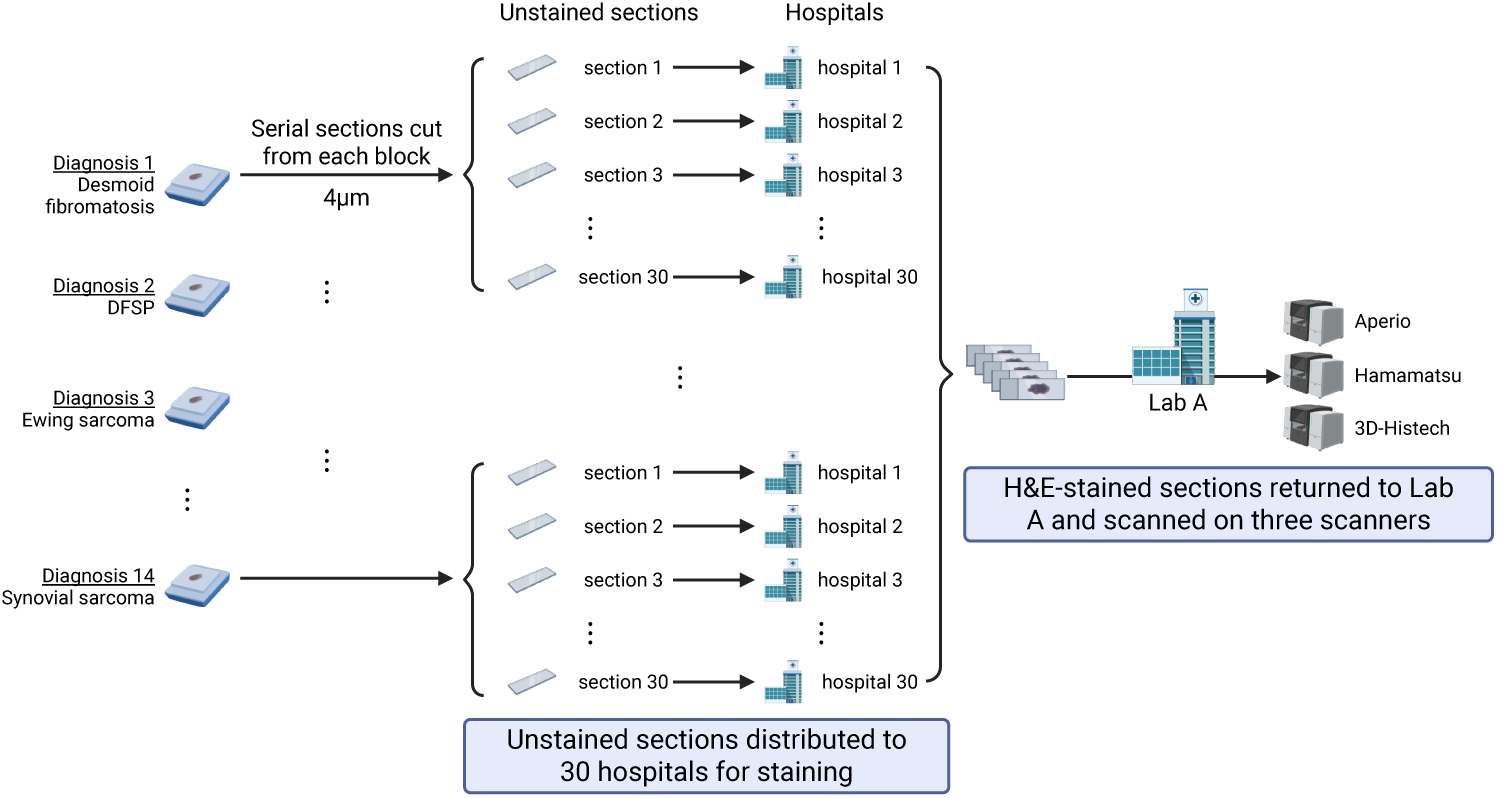
Preparation of a unique multi-institutional multi-scanner collection of slides (Cohort 2, n = 1, 243 WSIs). This collection comprises a representative case from each of the 14 tumour subtypes, from which 30 serial sections (4µm) were cut from one block and sent to different National Health Service (NHS) laboratories for H&E staining according to the local team’s standard operating procedures. Stained slides were then returned to Lab A and scanned on three different scanners (Leica Aperio GT 450, Hamamatsu NanoZoomer S360 and 3D-Histech Pannoramic 1000). DFSP, dermatofibrosarcoma protuberans.

### 2.3 How robust are foundation models to scanning variation?

We next investigated how variability in colour induced by different scanning devices affects the generalisability of foundation models. Given our limited availability of non-Aperio scanned slides, we aimed to isolate potential performance degradation introduced by mismatched scanner types during training (Fig. 5 **a**). For this experiment, we built a cohort (Cohort 3, Table 0) comprising eight cases randomly selected from each of the 14 diagnoses, stained at a single institution (Lab A) and scanned on three scanners. These were used for training three classifiers for each foundation model: an Aperio-trained model, a Hamamatsu-trained model and a 3D-Histech-trained model (Fig. 5 **a**). Results (Fig. 5 **b**) showed that overall the models performed best when the training and testing WSIs were scanned using the same platform (Aperio). CONCH-v1.5, Virchow, PRISM, and ResNet50 exhibited a noticeable drop in classification accuracy when trained on slides scanned with Hamamatsu or 3D-Histech but tested on Aperio-scanned slides. In contrast, TITAN and UNI-v2 demonstrated greater consistent performance across scanner conditions: both remained high across all scanners, suggesting their robustness and generalisability to scanning variation.

**Fig. 4.**
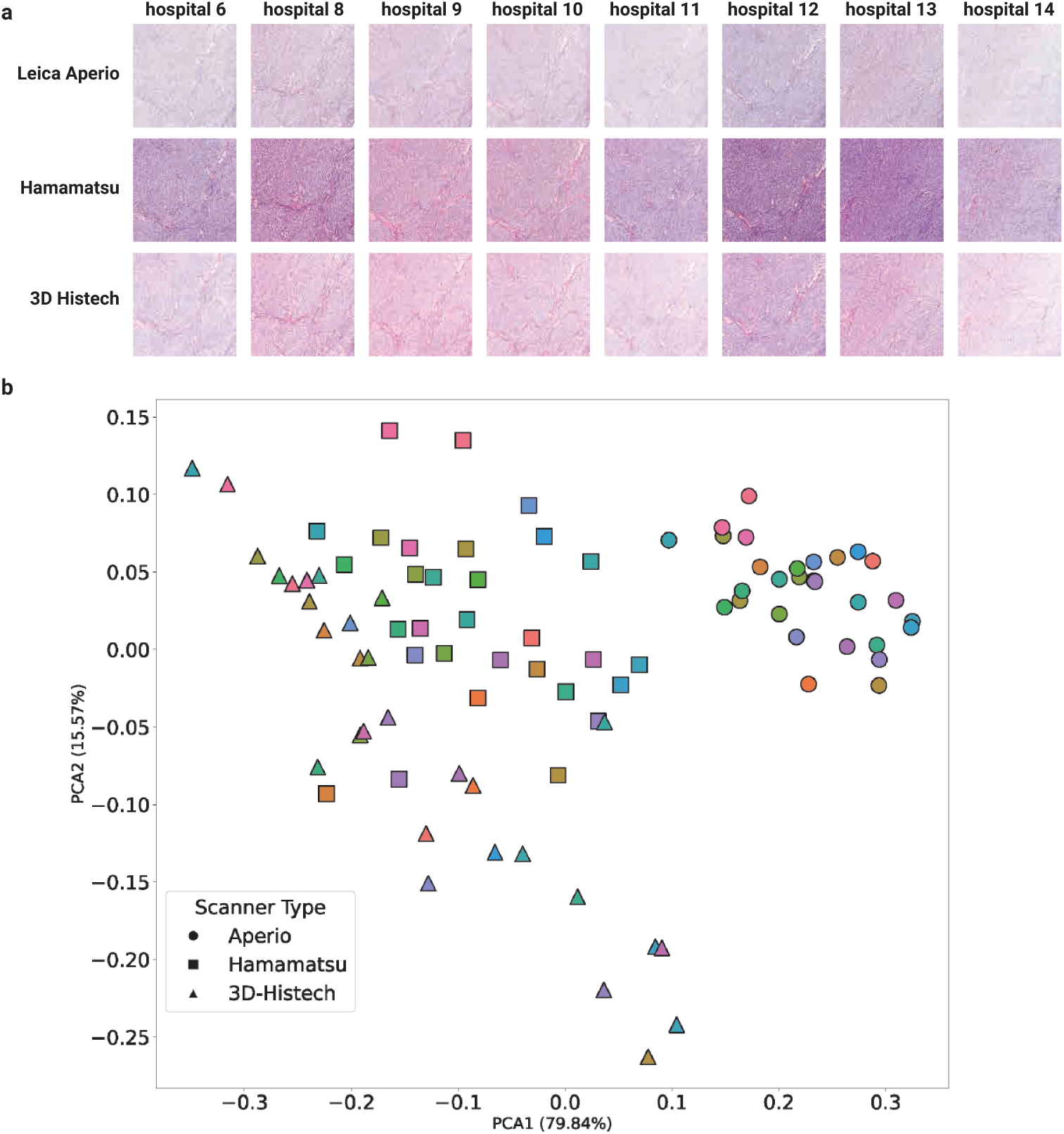
Visual and quantitative assessment of colour variation across staining institutes and scanners. a, Representative images of a synovial sarcoma stained at 8 (out of 30) laboratories and scanned using three different WSI (Leica Aperio GT 450, Hamamatsu NanoZoomer S360, and 3D-Histech Pannoramic 1000), 3000 × 3000 pixel tiles extracted. b, Principal component analysis (PCA) of stain vectors extracted from each 3000 × 3000 pixel tile. Each symbol represents a tile, where the colours correspond to the staining institute and the shape denotes the scanner used.

**Fig. 5.**
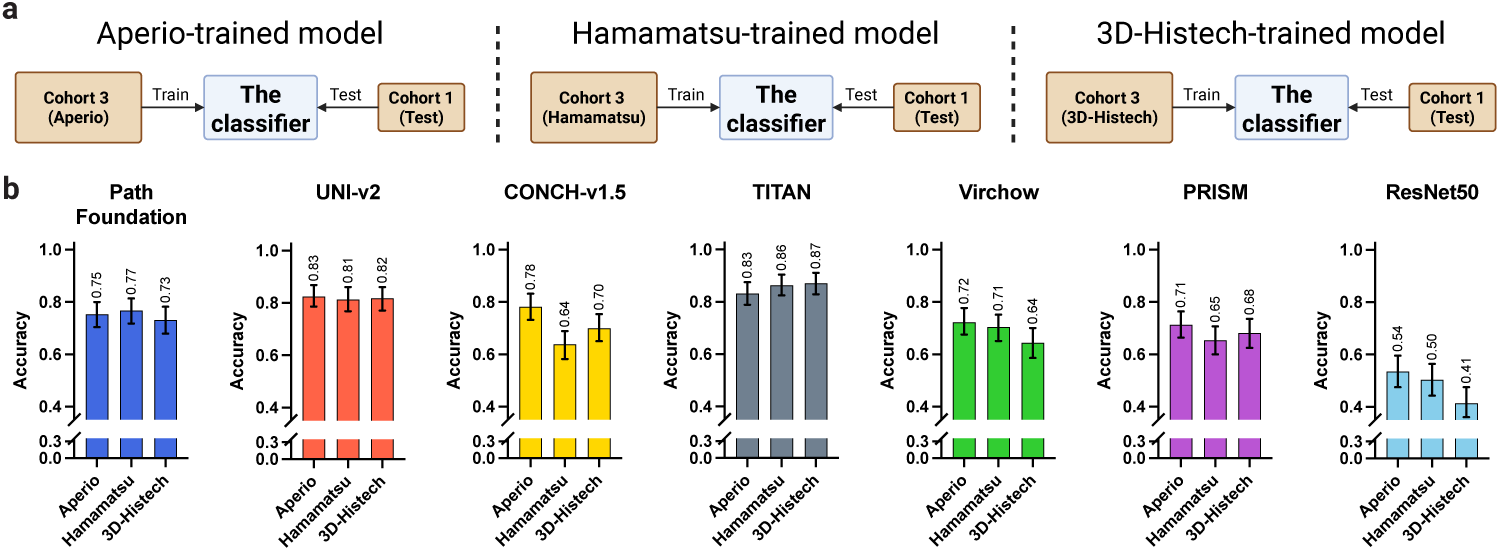
Controlled scan variation experiments. a, Schematic overview of the controlled scan variation experiment. A cohort of eight cases per diagnosis (Cohort 3), all stained at Lab A, was scanned on three different devices. Three separate classifiers were trained on identical sets of slides scanned with different scanners (Aperio, Hamamatsu and 3D-Histech scanners, respectively) and then evaluated on the same test set (Cohort 1-Test, stained at RONH, scanned with Aperio). b, Results of the controlled scan variation experiment for seven models.

### 2.4 Do foundation models enable few-shot learning for rare tumour classification under stain variation?

To evaluate the data efficiency of foundation models in the context of rare tumours, and to assess their robustness to stain variation under limited data, we simulated low-data settings using Cohort 3. Classifiers were trained using a small number of slides from each scanner type and evaluated on Cohort 1-Test. Few-shot training was conducted with increasing numbers of slides per diagnosis (1, 2, 4, 6, and 8).

TITAN demonstrated strong performance even when trained with one slide per diagnosis, while other models struggled with one slide for training (Fig. 6). However, patch-based encoders such as UNIv2 caught up quickly as the number of training slides increased. All foundation models outperformed ResNet50 in the few-shot setting. All models perform better when trained with the slides scanned using the same scanner as the test slides (Aperio, Fig. 6 **a**). However, similar improved performance patterns were observed regardless of the scanner of the training slides.

**Fig. 6.**
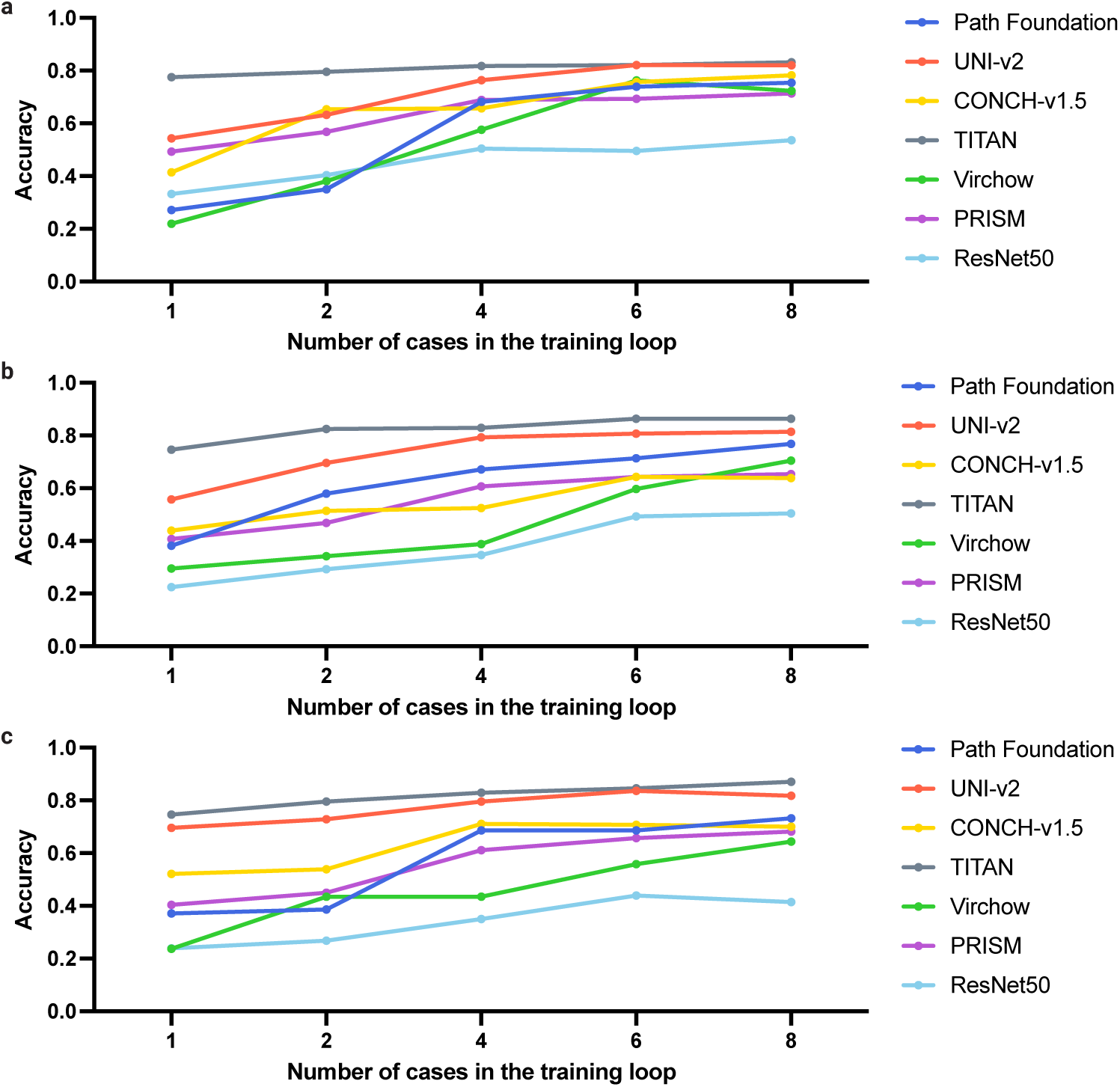
Sample size sensitivity analysis using single-institutionally stained, multiple-device scanned slides, they are: Lab A-stained Aperio-scanned (a), Lab A-stained Hamamatsu-scanned (b), and Lab A-stained 3D-Histech-scanned (c). Classifiers were trained using 1, 2, 4, 6, and 8 case(s) per diagnosis, and then evaluated on the same held-out test set. Despite limited training data, most foundation models achieved strong performance (over 0.8 accuracy) with as few as 8 cases per subtype, highlighting their sample efficiency in rare disease settings.

## 3 Discussion

We systematically benchmarked seven deep learning models, including ResNet50, Google Path Foundation, UNI-v2, CONCH-v1.5, TITAN, Virchow, and PRISM, on 14 different types of soft tissue tumours. These models span conventional convolutional neural networks to recent foundation models, with TITAN and PRISM representing slide-level encoders. Foundation models consistently outperformed traditional CNNs in both internal and external evaluations, demonstrating greater robustness to interinstitutional variation. However, even the best-performing models showed a 6–10% drop in accuracy on multi-institutionally stained test sets. This performance gap was substantially reduced by incorporating a small number of externally stained slides into training the classifier, underscoring the value of including representative staining variation to improve generalisability. Additionally, we identified scanner-related variation as a dominant source of performance degradation. Overall, while foundation models are not fully invariant to staining and scanning differences, fine-tuned classifiers based on foundation models are strong candidates for real-world pathology applications.

This study focuses on soft tissue tumor classification, but we have used these diagnoses as a proxy task to study stain variation and foundation models, and the findings and methodology presented here are broadly applicable to other types of tumor and computational pathological tasks. Sarcomas, a heterogeneous group of mesenchymal malignancies, represent less than 2% of all cancers [33]. They comprise numerous subtypes that often share overlapping histological features with benign lesions [37], making accurate diagnosis challenging, which can lead to delays in the diagnostic pathway [33, 38]. Given these challenges, patients with these tumours or their mimics could benefit from AI-powered Clinical Diagnostic Support Systems. Multi-instance learning (MIL) frameworks and self-supervised pretraining algorithms [6, 39, 40] could be exploited to address intra-tumoural heterogeneity and data scarcity.

Another key finding of this study is the strong data efficiency of the slide encoder TITAN. Unlike common cancers, where large training datasets are often available, sarcoma highlights the need for data-efficient learning. In few-shot experiments using only 1 to 8 cases per diagnosis, TITAN maintained stable performance, achieving over 0.75 accuracy even with a single case per class. This demonstrates TITAN’s strong discriminative power in rare tumour settings, where large, well-annotated datasets are difficult to obtain. These results underscore the potential of employing slide-level foundation models for classifying rare diseases such as sarcoma. Nevertheless, it is noteworthy that as the training set size increased, patch-based encoders showed rapid performance gains, suggesting that while slide encoders excel in low-data regimes, patch encoders may match or surpass them when data are abundant.

To advance research in this field, we are releasing a unique multi-institutional dataset comprising over 1,200 whole slide images covering 14 tumour types, stained in 30 institutions, and scanned with three scanners (Fig. 3). This resource could facilitate further research into the generalisability of AI models in digital pathology, particularly in the context of staining and scanning variations.

This study has several limitations. First, only three scanner types (Aperio, Hamamatsu, and 3D-Histech) were evaluated. While we believe these scanners represent common real-world variation, the inclusion of additional devices, such as those from Philips and Roche, could further validate the generalisability of our findings. Second, we focused on a single task: soft-tissue tumour classification; future work could expand this evaluation on stain variations to additional tumour types and tasks, such as mitosis detection and genomic marker prediction [41, 42].

Several avenues remain open for further exploration. Current stain normalisation and augmentation strategies are primarily image-based and were originally developed for conventional CNNs. There is a notable lack of embedding-space augmentation or normalisation techniques specifically tailored to foundation models, which is an open area with significant potential. Developing such methods could further stabilise downstream performance across diverse histological and technical conditions.

In summary, this study leverages sarcomas and their histological mimics as a testbed to evaluate the robustness of foundation models to real-world variations in staining and scanning. Through benchmarking and controlled experiments, we demonstrate that foundation models exhibit promising resilience to such variability, when even a small number of training samples reflecting these differences are included. By releasing a curated, multi-institutional dataset, we aim to support the development of more generalisable and robust AI models for digital pathology. While foundation models show significant promise, generalisation to external data remains a major challenge in building reliable AI-based diagnostic systems. Increasing the diversity of training data continues to be a simple yet effective strategy for improving model generalisability.

## 4 Methods

**Ethical approval declarations** Ethical approval was given for undertaking the study ’An Artificial Intelligence (AI) solution for diagnosing, prognosticating as well as predicting the outcome of sarcomas and their mimics: a multi-centre study’. IRAS project ID: 328987 Protocol number: EDGE 161548. REC reference: 23/NI/0166. Sponsor University College London has been approved by HRA and Health and Care Research Wales (HCRW) (December 14^th^ 2023) and by Health and Social Care Research Ethics Committee B (HSC REC B) Office for Research Ethics Committees Northern Ireland (ORECNI) Lissue Industrial Estate West, 5 Rathdown Walk, LISBURN, BT28 2RF. REC reference: 23/NI/0166, Protocol number: EDGE 161548, IRAS project ID: 328987 (December 2023).

### 4.1 WSI datasets

We constructed four collections of hematoxylin and eosin (H&E)-stained WSI comprising 14 different diagnoses of sarcoma and their mimics. In total, 2,350 different WSIs were included in the study; each WSI was selected from a different case and only one image per patient. Table 0 displays the four collections. Diagnoses were based on the 2020 edition of the WHO series of soft tissue and bone tumour classification [33]. The tumours studied included desmoid fibromatosis, dermatofibrosarcoma protuberans (DFSP), Ewing sarcoma, glomus tumour, intramuscular myxoma, leiomyosarcoma, low-grade fibromyxoid sarcoma (LGFMS), lymphoma, myxoid liposarcoma, neurofibroma, nodular fasciitis, schwannoma, solitary fibrous tumour, and synovial sarcoma (Fig. 1 **b**). Each case was reviewed by at least three expert soft tissue tumour pathologists, and the diagnoses were established using state-of-the-art immunohistochemistry, molecular profiling and clinical context. The cohorts described below are found in Table 0 and all cases are listed in the supplementary spreadsheet.

**Cohort 1** serves as a controlled baseline for training and evaluation. It comprises 1,394 H&E-stained WSI representing approximately 100 different cases per 14 diagnoses (one case per patient). All cases were sourced, processed, and stained at Lab A, and subsequently digitised using Leica Aperio GT 450 scanners at 40× equivalent magnification (0.2629 *µ*m pixel*^−^*^1^). The WSIs were split into training (60%), validation (20%), and hold-out test (20%) sets. Lab A is UKAS-accredited [43], and all tasks are undertaken according to standard operating procedures.

**Cohort 2** was designed to investigate staining and scanning variability. See Fig. 3 for details. It was made up of 1,243 WSI which were different to those cases in Cohort

1. For each of the 14 tumour subtypes studied, a case with representative histology for each was selected and serial sections (4 *µ*m) were cut from a single tissue block. One unstained slide per case was distributed to 30 different National Health Service (NHS) hospital laboratories, excluding Lab A (see list of members of Consortium), where local teams performed H&E staining using their clinical laboratory protocols. The stained slides were returned and digitised using three slide scanners: Leica Aperio GT 450, Hamamatsu NanoZoomer S360, and 3D-Histech Pannoramic 1000 (Fig. 3).

**Cohort 3** was designed to assess the impact of scanning variation while controlling for staining variability (*n* = 335 WSIs). For this cohort, eight cases per diagnosis were selected from Cohort 1 or Cohort 2 apart from eight new cases of synovial sarcoma which were added. All cases were stained at Lab A. Each stained slide was then scanned using the three scanners: Leica Aperio GT 450, Hamamatsu Nanozoomer S360, and 3D-Histech Pannoramic 1000. One WSI was not available from the 3D-Histech batch. **Cohort 4** included all 14 diagnoses and serves as a ‘real-world’ test set. 934 WSIs were sourced from Lab A’s clinical archives; 105 were stained at Lab A and the remaining 838 were stained in 73 different hospitals. These slides are different to those in Cohort 1, 2 and 3; details are found in Extended Table B2). This cohort was divided into two groups; 813 slides were used exclusively to train Cohort 4 and the remaining 121 slides were used in combination with Cohort 1-train to form a Combined cohort for the purpose of investigating if training on a limited number of images stained in different institutions improvded the accuracy of the models.

### 4.2 Slides tiling

TRIDENT [35, 36] was used to preprocess the WSI (Fig. 1 **a**), during which all background tiles were efficiently discarded. Tiles can be generated under different magnification levels, at different sizes and with various levels of overlap between adjacent tiles. We used the default tiling settings for each of the models, which are: 224*224 at 20x for Path Foundation, UNI-v2, CONCH-v1.5, Virchow, PRISM and ResNet50, and 512*512 at 20x for TITAN.

### 4.3 Foundation models

We benchmarked the effectiveness of six foundation models for the sarcoma/sarcoma mimics subtyping task with staining and scanning variation. Path Foundation (Google Research) uses ViT-S/B backbone [44] trained with Masked Siamese Networks [45] on 60 million H&E tiles from the Cancer Genome Atlas (TCGA) dataset [8], sampled at multiple magnifications to promote resolution robustness. UNI-v2, CONCH-v1.5 and TITAN (all from Mahmood Lab) are also ViT-based models leveraging multiobjective [11], contrastive learning [12] and end-to-end slide-level pretraining [13], respectively. Virchow and PRISM (both from PaigeAI) are large-scale models trained on over 1.5 million WSIs across diverse tissues [9, 14]; PRISM uniquely integrates tile embeddings with clinical text using a multimodal generative approach. A ResNet50CNN [34], used as a benchmark model, was trained in a supervised manner directly on WSIs, without pretraining on external histopathology datasets or using self-supervised learning. This provided a reference point on which to evaluate the benefits of using foundation model-based feature extraction approaches in the presence of staining and scanning variation. Models were used either as tile encoders with Multi-Instance Learning (MIL) or as end-to-end slide classifiers.

### 4.4 Slide-level classification from tile- and slide-based embeddings

To derive slide-level diagnosis predictions, we adopted two strategies depending on the granularity of the feature representations.

For models producing patch-level embeddings (Google Path Foundation, UNI-v2, CONCH-v1.5, Virchow and ResNet50-CNN), we adopted an attention-based multi-instance learning (MIL) framework [40]. Each slide was treated as a bag of instances (i.e., tiles/patches), and the MIL model learned to assign an attention weight to each tile, reflecting its relative importance for the slide-level classification task. This attention mechanism could get those non-informative or noisy tiles naturally filtered out and enable the classification model to focus on diagnostically relevant information.

These weights are used to aggregate tile embeddings into a single slide-level representation via a weighted sum (*z* = ^L-^*^K^α_i_x_i_*). We randomly sampled *K* = 500 tiles per slide during training, or used all tiles if fewer were available. Models were trained with cross-entropy loss using the Adam optimiser and early stopping based on validation loss.

For slide-level encoders such as TITAN and PRISM, which directly output a fixed-length embedding per whole slide, we trained a logistic regression classifier using these slide-level vectors. The classifier was implemented with scikit-learn [46], trained on the slide embeddings and corresponding labels without regularisation or hyperparameter tuning. A maximum of 1000 iterations and a fixed random seed ensured convergence and reproducibility.

### 4.5 Statistical analysis

To evaluate differences in classification performance, Pearson’s chi-squared test of independence was applied in two settings: (i) to assess whether each model’s classification accuracy differed significantly between the internal (Cohort 1-Test) and multi-institutional (Cohort 4-Test) test sets, and (ii) to test whether if a small amount of multi-institutional slides in training (i.e. trained with the Combined cohort, Fig. 2 **a**) significantly improved performance over training with only the internal slides (i.e., Cohort 1). For each comparison, the number of correctly and incorrectly classified slides was tabulated and used to compute the test statistic. All tests were two-sided, with significance defined as *p <* 0.05.

To quantify the uncertainty around classification performance, we estimated 95% confidence intervals for the accuracy using non-parametric bootstrapping. Specifically, we resampled the test set with replacement 1,000 times, preserving its original size in each iteration. For each resampled set, accuracy was recalculated, yielding a distribution of accuracy estimates. The 2.5*^th^* and 97.5*^th^* percentiles of this distribution were taken as the lower and upper bounds of the confidence interval. This approach makes no assumptions about the underlying distribution of accuracy and provides a robust, data-driven measure of variability.

### 4.6 Model evaluation

For each experiment, model performance was evaluated on the held-out test set using standard metrics. These included micro-averaged accuracy (i.e., the overall proportion of correctly predicted diagnoses across all test slides), top-k accuracies (where *k* = 1, 3) and class-wise (i.e., subtype-level) accuracy breakdowns. To support interpretability and clinical transparency, each classifier also provided the top-3 predicted classes along with their associated confidence scores for every test slide. These results were exported as structured CSV reports to facilitate downstream analysis and visualisation. This evaluation framework enables a lightweight and interpretable classifier to be applied on top of the rich representations extracted by foundation models, requiring significantly less annotated data and computational resources than training an end-to-end deep neural network.

## Data availability

A unique multi-institutional dataset comprising 1,243 whole slide images covering 14 tumour types, stained in 30 external institutions in addition to Laboratory A where the project was carried out, and scanned with three scanners is shared with the research community at: LINK. We provide an interactive tool for (1) Fig. 1 **d**, **e**, and (2) Fig. A2 for the exploration of the qualitative t-SNE outputs, at: https://cbhindex.github.io/visualise_scan_stain_efffects.

## Code availability

Python implementation of the methodology and documentation are at: https://github.com/cbhindex/path_foundation_variation.

## Consortia collaborators

The members of the AI Scope Consortium involved in this project are: Adrienne Flanagan (UCL and Royal National Orthopaedic Hospital), Anna Tollit (UCL and Royal National Orthopaedic Hospital), Fatine Oumlil (UCL and Royal National Orthopaedic Hospital), Fernanda Amary (UCL and Royal National Orthopaedic Hospital), Roberto Tirabosco (UCL and Royal National Orthopaedic Hospital), Vahghelita Andrei (UCL and Royal National Orthopaedic Hospital), Nischalan Pillay (UCL and Royal National Orthopaedic Hospital), Paul Cool (The Robert Jones and Agnes Hunt Orthopaedic Hospital), Adam Levine (UCL and Royal Free London NHS Foundation Trust), Sebastian Brandner (UCL and University College London Hospital NHS Foundation Trust), Angela Richard-Londt (UCL and University College London Hospital NHS Foundation Trust), Petra Balogh (University Hospitals Birmingham NHS Foundation Trust), Phillipe Taniere (University Hospitals Birmingham NHS Foundation Trust), Preethi Gopinath (Princess Alexandra Hospital NHS Trust), Neil Sebire (Great Ormond Street Hospital for Children NHS Foundation Trust and UCL), Luis Campos (Great Ormond Street Hospital for Children NHS Foundation Trust), Thomas Jacques (Great Ormond Street Hospital for Children NHS Foundation Trust and UCL), Sarah Coupland (Royal Liverpool University Hospital), Naomi Kamanga (Royal Liverpool University Hospital), Katalin Boros (Manchester University NHS Foundation Trust), Edmund Cheesman (Manchester University NHS Foundation Trust), Elizabeth Halloran (Manchester University NHS Foundation Trust), Caroline Glennie (Manchester University NHS Foundation Trust), Leona Doyle (University College Dublin, Ireland), Adrian Marino-Enriquez (University College Dublin, Ireland), Jonathan Davey (NHS Lothian), Hannah Monaghan (NHS Lothian), Michael Toss (Sheffield Teaching Hospitals NHS Foundation Trust), David Hughes (Sheffield Teaching Hospitals NHS Foundation Trust), Malee Fernando (Sheffield Teaching Hospitals NHS Foundation Trust), David Leff (Sheffield Teaching Hospitals NHS Foundation Trust), Patrick Shenjere (The Christie NHS Foundation Trust), Oisin Houghton (Belfast Health and Social Care Trust), Tom McCulloch (Nottingham University Hospitals NHS Trust), Filomena Medeiros (Basildon University Hospital, Mid and South Essex NHS Foundation Trust), Ann Sandison (Guy’s and St Thomas’ NHS Foundation Trust), Kalnisha Naidoo (King’s College Hospital NHS Foundation Trust), Getnet Demissie (King’s College Hospital NHS Foundation Trust), Heena Patel (King’s College Hospital NHS Foundation Trust), Uchechi Igbokwe (Queen’s Hospital, Barking, Havering and Redbridge University Hospitals), Graham Thwaites (Queen’s Hospital, Barking, Havering and Redbridge University Hospitals), Keeley Thwaites (Queen’s Hospital, Barking, Havering and Redbridge University Hospitals), Ann Fleming (Maidstone and Tunbridge Wells NHS Trust), Mercy Berin (Maidstone and Tunbridge Wells NHS Trust), Shyamala Fernandez (Ealing Hospital, London North West University Healthcare NHS Trust), Rajesh Nalluri (Northwick Park Hospital, London North West University Healthcare NHS Trust), Zahra Atiyyah (Northwick Park Hospital, London North West University Healthcare NHS Trust), Konrad Wolfe (Southend University Hospital, Mid and South Essex NHS Foundation Trust), Victoria Hills (Pathology First, SYNLAB UK & Ireland), Tomos Saunders (Pathology First, SYNLAB UK & Ireland), Matilda Ralph (Hemel Hempstead Hospital, West Hertfordshire Teaching Hospitals NHS Trust), Nicholas Southgate (Hemel Hempstead Hospital, West Hertfordshire Teaching Hospitals NHS Trust), Francesca Maggiani (North Bristol NHS Trust), Zsolt Orosz (Oxford University Hospitals NHS Trust), Jennifer Brown (Oxford University Hospitals NHS Trust), Nick Athanasou (Oxford University Hospitals NHS Trust), Ian Cook (Salisbury NHS Foundation Trust), Sarah Oliver (Salisbury NHS Foundation Trust), Peter Davis (Broomfield Hospital, Mid and South Essex NHS Foundation Trust), Shelley Brown (Broomfield Hospital, Mid and South Essex NHS Foundation Trust), Jasenka Mazibrada (Norfolk and Norwich University Hospitals NHS Foundation Trust), Nathan Asher (University College Hospital NHS Foundation Trust), Efren Quilala (Queen Elizabeth Hospital, Lewisham and Greenwich NHS Trust), Dalian Beaver (Queen Elizabeth Hospital, Lewisham and Greenwich NHS Trust), Izhar Bagwan (Royal Surrey County Hospital, Royal Surrey NHS Foundation Trust), Dawn Whyndham (Royal Surrey County Hospital, Royal Surrey NHS Foundation Trust), Khin Thway (Royal Marsden NHS Trust), Michael Hubank (Institute of Cancer Research), Janet Shipley (Institute of Cancer Research), and Anna Kelsey (Royal Manchester Children’s hospital)

## Acknowledgments

This project was supported by EPSRC UKRI Funding Services Grant Ref: EP/Y020030/, Sarcoma UK (SUKRI.2023), BCRT (BCRT/5717) and the Royal National Orthopaedic Hospital Research. AMF is supported by the National Institute for Health Research, UCLH Biomedical Research Centre, and the CRUK Experimental Cancer Centre. TC is supported by the UCLH Biomedical Research Centre and the Turing-Roche Strategic Partnership. We thank the UCL/UCLH Biobank for Studying Health and Disease for the provision of human tissue samples and clinical data and are grateful to the Biobank Team at the RNOH and all healthcare workers who cared for the patients without whose input this work would not have been possible.

## Appendix A Extended Data: Embedding space exploration by visualising the features in two dimensions

**Fig. A1.**
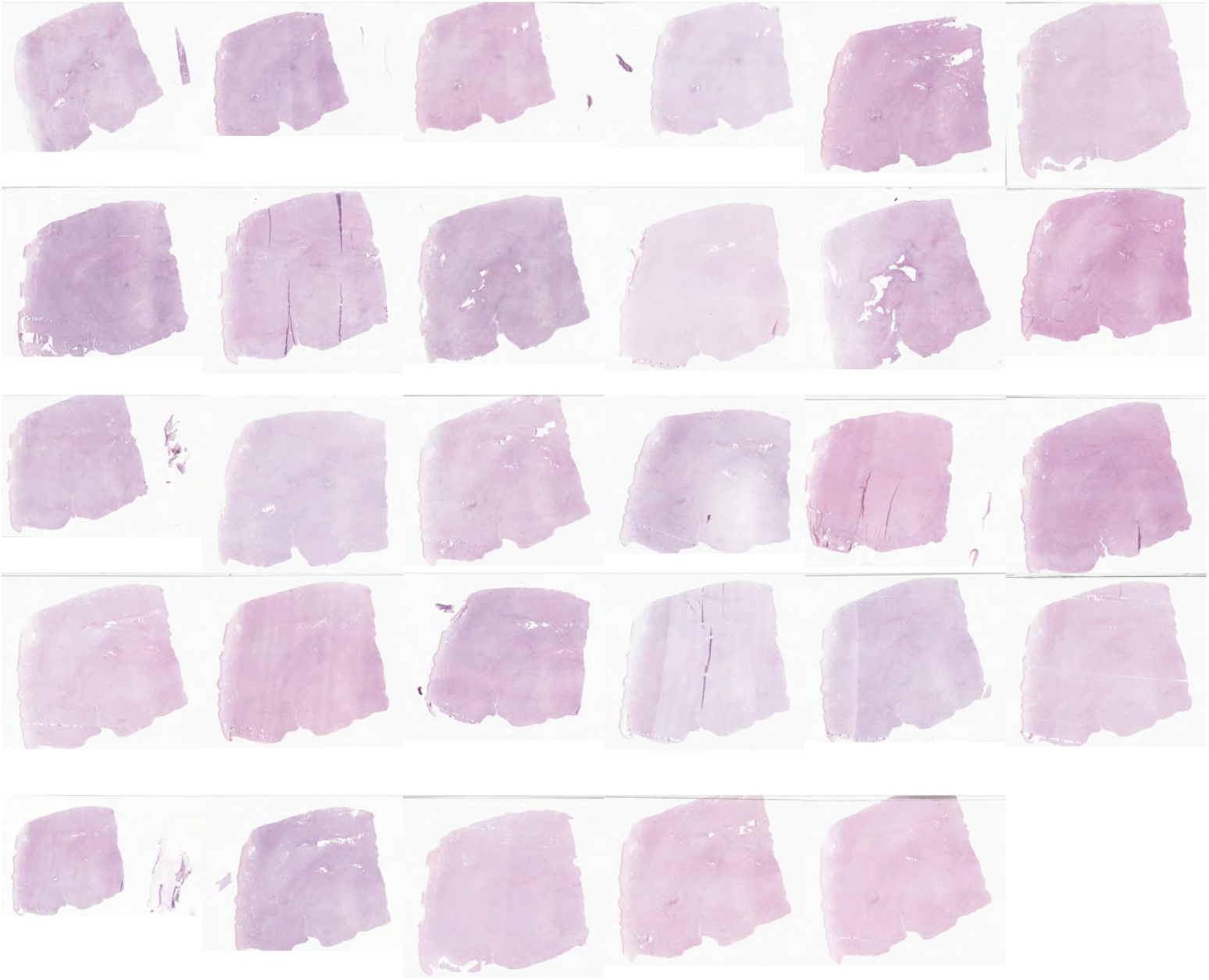
Serial sections from a sample of a leiomyosarcoma stained at 30 different NHS Trusts show significant colour variation after digitisation with the same Aperio scanner. One slide was not returned.

One of the most visually apparent challenges in real-world digital pathology is the substantial colour variation introduced by differing staining protocols across institutions. As illustrated in Fig.A1, serial sections from a single leiomyosarcoma case were circulated to multiple NHS Trusts, each of which applied their routine haematoxylin and eosin (H&E) staining procedures. Despite being scanned using the same Aperio scanner under controlled imaging conditions, the resulting slides exhibited considerable variation in hue, contrast, and saturation. This example underscores the difficulty of achieving colour consistency in multi-centre datasets and highlights the necessity for developing robust AI systems that can generalise across diverse staining conditions.

To qualitatively assess the types of staining and scanning information captured by foundation models, we visualised the feature/embedding space using two-dimensional t-distributed Stochastic Neighbour Embedding (t-SNE) for the staining- and scanning-controlled dataset (Cohort 2, Table 0 and Fig. 3). The projection was performed on the two best-performing models, UNI-v2 and TITAN, using tile-level embeddings and slide-level embeddings, respectively. As UNI-v2 is a tile-level encoder, we randomly sampled three informative tiles (i.e., tiles from tumour areas) per slide and employed their tile-level embeddings for the t-SNE plot. This setting ensures consistent comparison across slides. In contrast, TITAN has a two-stage architecture, containing a tile-level encoder followed by a slide-level aggregator, so a single slide-level embedding vector was obtained per whole slide image. These slide-level embeddings were employed to represent an entire slide on the t-SNE plot. To facilitate interpretability, the dots in each of the t-SNE plots were coloured by three distinct categorical schemes:

(1) the diagnostic label as the slide-level ground truth, (2) scan device, and (3) staining institute (Fig. A2). These embedding space plots offered insights into how well the learned representations disentangle the relative contributions of biological, staining and scanning factors to model robustness and generalisability. An interactive tool, implemented with scikit-learn [46] and Plotly [47], is provided to explore the t-SNE outputs (see Data Availability).

Both UNI-v2 and TITAN demonstrated strong class-specific clustering when dots in t-SNE plots were coloured by the slide-level diagnosis: most tumour subtypes forming compact and well-separated groups (Fig. A2 **a**, **b**). This pattern indicates that the embeddings encode meaningful biological information that reflects histological differences between tumour types. Even morphologically similar subtypes, such as lymphoma and Ewing sarcoma, appeared as distinct clusters, indicating that both models are sensitive to diagnostically relevant features. These results align with the strong subtype classification performance observed in earlier sections (Fig. 1 **c**, Fig. 2 **b** and Table B3), reinforcing the view that foundation models offer robust feature extraction capabilities even in complex and heterogeneous diseases like sarcoma. Notably, the clustering observed in TITAN was more distinct and less noisy than that of UNI-v2. The TITAN embeddings produced clearly delineated diagnostic groups with minimal overlap (Fig. A2 **b**), reflecting the strength of its two-stage architecture to a certain degree, particularly the slide-level encoder. In contrast, while UNI-v2 also exhibited meaningful clustering, some cases appeared more dispersed or intermixed (Fig. A2 **a**), which may stem from the fact that only three random tiles were sampled per slide. These sampled tiles may not always represent tumour-rich regions, thereby introducing variability into the embedding space. Overall, these results highlight the added benefit of slide-level modelling in enhancing representation quality and diagnostic separability. When the same embeddings were coloured by scanner device, a different pattern emerged between the two models. In the UNI-v2 t-SNE plot (Fig. A2 **c**), three dominant sub-clusters became visible within each diagnostic group, aligning with the scanner used. This suggests that despite capturing biologically meaningful features, the tile-level embeddings from UNI-v2 remain moderately sensitive to the image acquisition device. In contrast, the TITAN embeddings (Fig. A2 **d**) exhibited minimal separation by scanner, with diagnostic clusters forming coherent and compact groupings regardless of the scanning device. This indicates that TITAN’s slide-level encoder effectively mitigates scanner-related artefacts in the embedding space. By contrast, when the embeddings were coloured by the staining institute (Fig. A2 **e**, **f** ), neither model showed any consistent or interpretable clustering. Points from different centres were broadly interspersed without discernible grouping, suggesting that both models are comparatively robust to staining protocol variation across institutions.

These findings suggest that foundation models such as UNI-v2 and TITAN prioritise biological signal over staining or scanning variations. While some scanner-related structure remains in tile-level models, slide-level representations, such as those from TITAN, achieve greater invariance. This further supports the utility of foundation models, particularly those incorporating whole-slide context, as robust and transferable embedding generators for multi-institutional digital pathology applications.

**Fig. A2.**
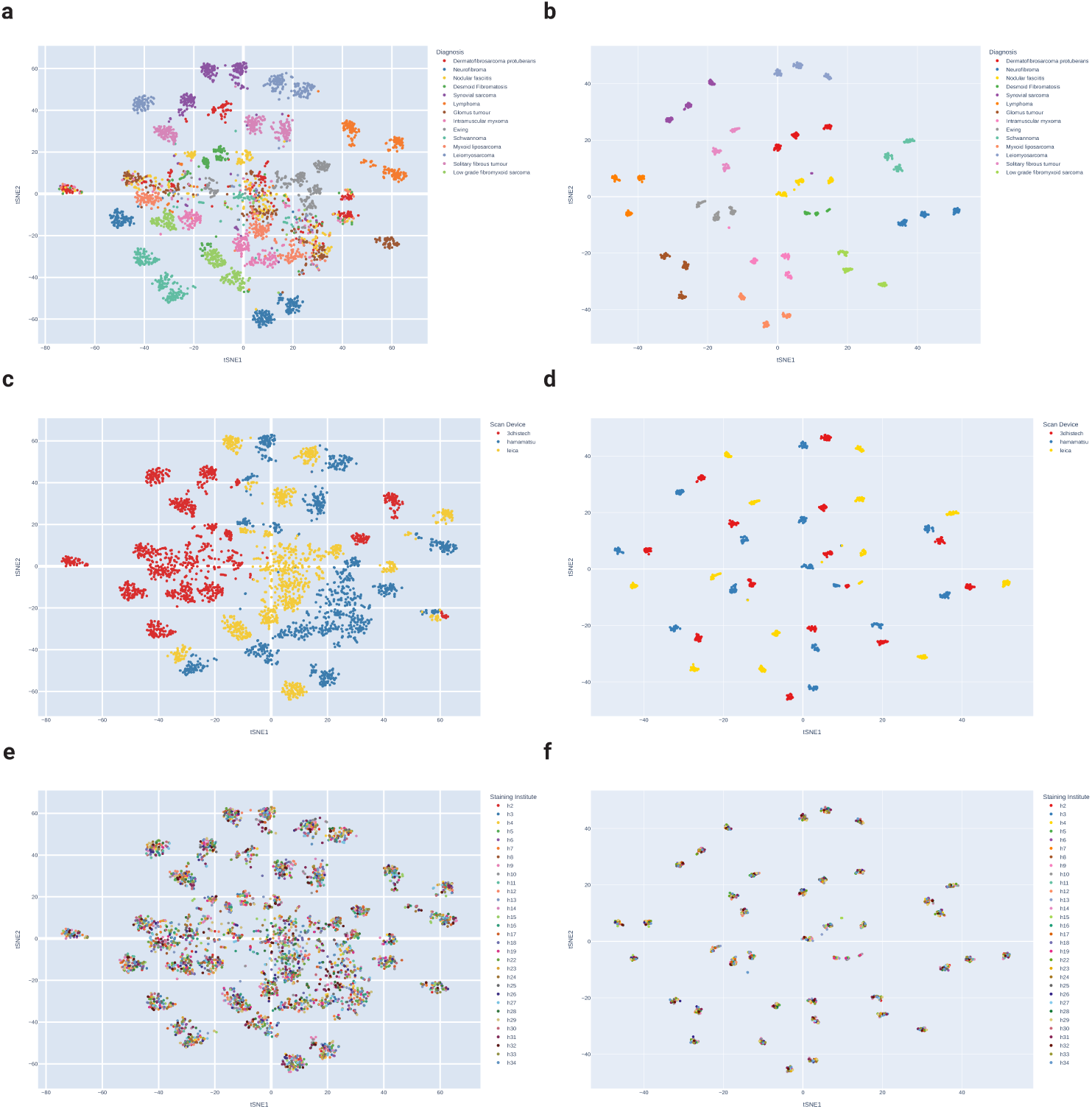
Visualisation of the feature space using t-SNE. a, c, e, Tile-level embeddings from the UNI-v2 model for the staining- and scanning-controlled dataset (Cohort 2). For each slide, three tiles were randomly sampled, resulting in three points per slide in the visualisation. The embeddings are coloured by diagnosis (a), scan device (c), and staining institute (e), respectively. b, d, f, Slide-level embeddings from the TITAN model for the same dataset, with one point per slide, coloured by diagnosis (b), scan device (d), and staining institute (f ). An interactive version of these t-SNE plots is provided (see Data Availability section).

## Appendix B Extended Tables

**Table B1.** Top-1 and top-3 classification accuracies on Cohort 4-Test set for ‘trained with Cohort 1’ and ‘trained with the Combined cohort’.

|  | Google | UNI-v2 | CONCH-v1.5 | TITAN | Virchow | PRISM | ResNet50 |
| --- | --- | --- | --- | --- | --- | --- | --- |
| **Top-1 (trained with Cohort 1)** | 0.721 | 0.847 | 0.772 | 0.860 | 0.852 | 0.782 | 0.533 |
| **Top-1 (trained with the Combined cohort)** | 0.808 | 0.899 | 0.838 | 0.881 | 0.865 | 0.803 | 0.736 |
| **Top-3 (trained with Cohort 1)** | 0.934 | 0.970 | 0.938 | 0.968 | 0.963 | 0.940 | 0.844 |
| **Top-3 (trained with the Combined cohort)** | 0.946 | 0.983 | 0.970 | 0.970 | 0.967 | 0.958 | 0.923 |

Footnote: All models exhibited substantial gains when training on the Combined cohort.

**Table B2.**
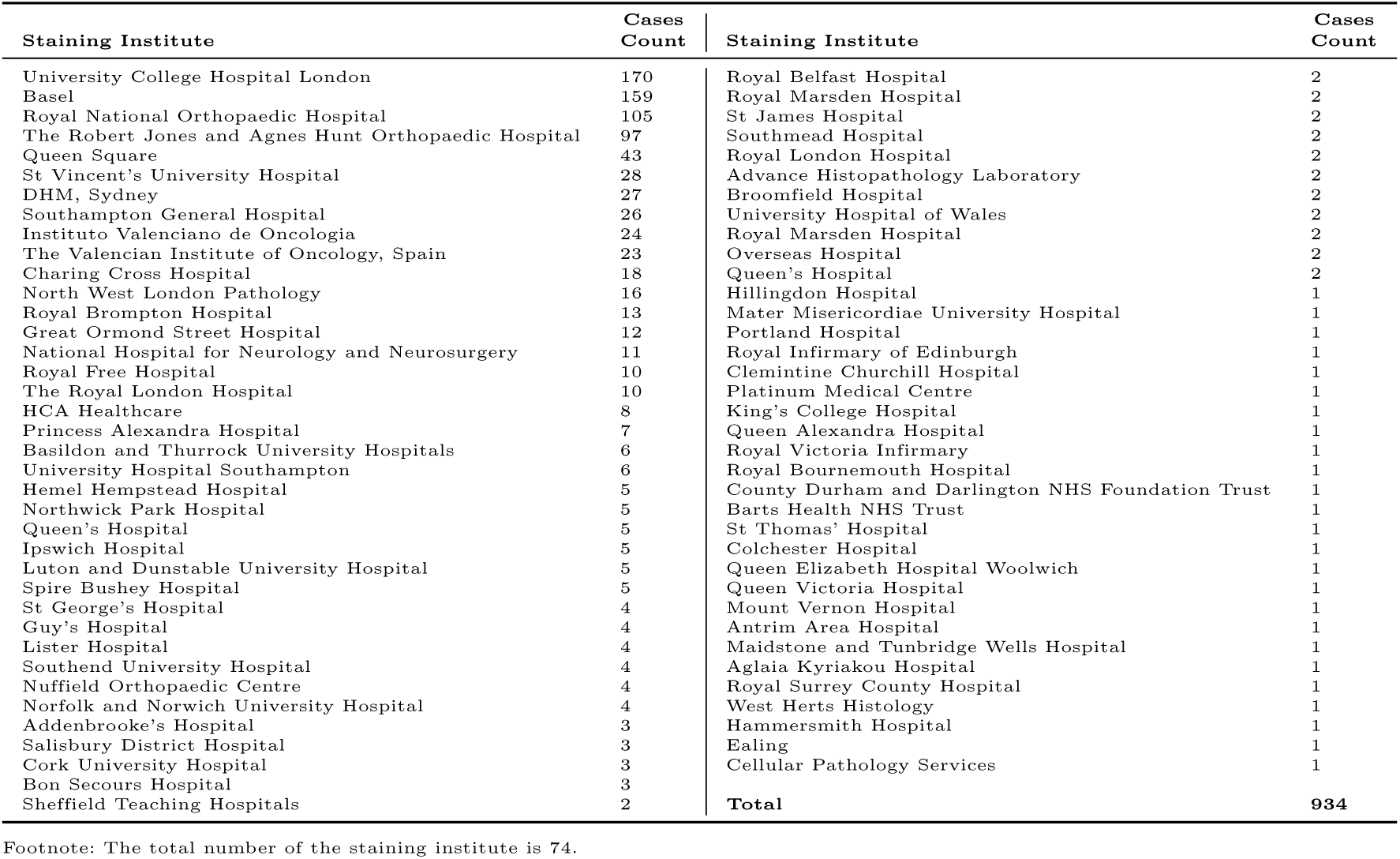
Cohort 4 cases distribution.

**Table B3.**
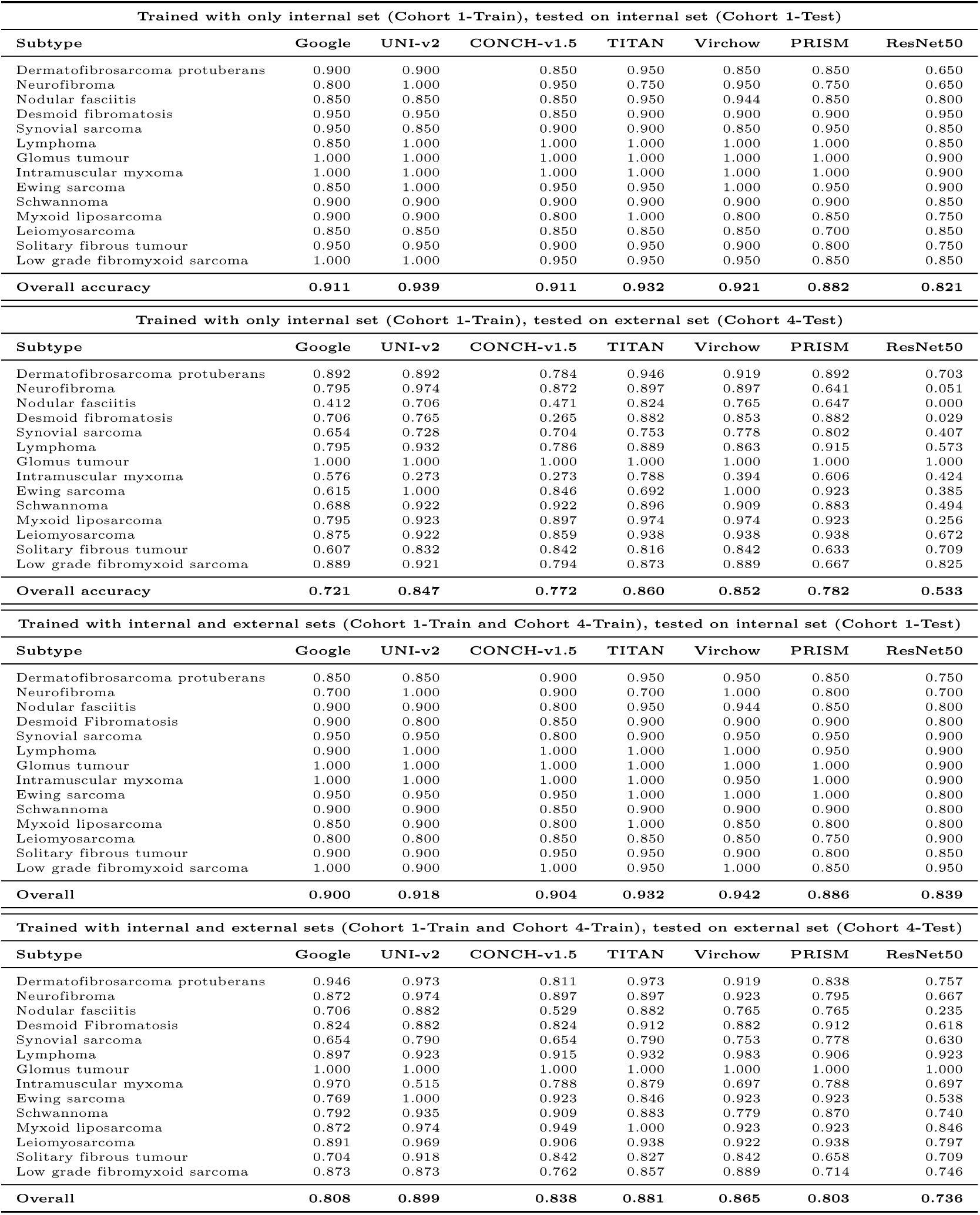
Classwise accuracies for soft tissue tumours across internal and multi-institutional test sets, using seven models: Google Path Foundation, UNI-v2, CONCH-v1.5, TITAN, Virchow, PRISM, and ResNet50-CNN.

| Trained with only internal set (Cohort 1-Train), tested on internal set (Cohort 1-Test) |  |  |  |  |  |  |  |
| --- | --- | --- | --- | --- | --- | --- | --- |
| Subtype | Google | UNI-v2 | CONCH-v1.5 | TITAN | Virchow | PRISM | ResNet50 |
| Dermatofibrosarcoma protuberans | 0.900 | 0.900 | 0.850 | 0.950 | 0.850 | 0.850 | 0.650 |
| Neurofibroma | 0.800 | 1.000 | 0.950 | 0.750 | 0.950 | 0.750 | 0.650 |
| Nodular fasciitis | 0.850 | 0.850 | 0.850 | 0.950 | 0.944 | 0.850 | 0.800 |
| Desmoid fibromatosis | 0.950 | 0.950 | 0.850 | 0.900 | 0.900 | 0.900 | 0.950 |
| Synovial sarcoma | 0.950 | 0.850 | 0.900 | 0.900 | 0.850 | 0.950 | 0.850 |
| Lymphoma | 0.850 | 1.000 | 1.000 | 1.000 | 1.000 | 1.000 | 0.850 |
| Glomus tumour | 1.000 | 1.000 | 1.000 | 1.000 | 1.000 | 1.000 | 0.900 |
| Intramuscular myxoma | 1.000 | 1.000 | 1.000 | 1.000 | 1.000 | 1.000 | 0.900 |
| Ewing sarcoma | 0.850 | 1.000 | 0.950 | 0.950 | 1.000 | 0.950 | 0.900 |
| Schwannoma | 0.900 | 0.900 | 0.900 | 0.900 | 0.900 | 0.900 | 0.850 |
| Myxoid liposarcoma | 0.900 | 0.900 | 0.800 | 1.000 | 0.800 | 0.850 | 0.750 |
| Leiomyosarcoma | 0.850 | 0.850 | 0.850 | 0.850 | 0.850 | 0.700 | 0.850 |
| Solitary fibrous tumour | 0.950 | 0.950 | 0.900 | 0.950 | 0.900 | 0.800 | 0.750 |
| Low grade fibromyxoid sarcoma | 1.000 | 1.000 | 0.950 | 0.950 | 0.950 | 0.850 | 0.850 |
| **Overall accuracy** | **0.911** | **0.939** | **0.911** | **0.932** | **0.921** | **0.882** | **0.821** |

| Trained with only internal set (Cohort 1-Train), tested on external set (Cohort 4-Test) |  |  |  |  |  |  |  |
| --- | --- | --- | --- | --- | --- | --- | --- |
| Subtype | Google | UNI-v2 | CONCH-v1.5 | TITAN | Virchow | PRISM | ResNet50 |
| Dermatofibrosarcoma protuberans | 0.892 | 0.892 | 0.784 | 0.946 | 0.919 | 0.892 | 0.703 |
| Neurofibroma | 0.795 | 0.974 | 0.872 | 0.897 | 0.897 | 0.641 | 0.051 |
| Nodular fasciitis | 0.412 | 0.706 | 0.471 | 0.824 | 0.765 | 0.647 | 0.000 |
| Desmoid fibromatosis | 0.706 | 0.765 | 0.265 | 0.882 | 0.853 | 0.882 | 0.029 |
| Synovial sarcoma | 0.654 | 0.728 | 0.704 | 0.753 | 0.778 | 0.802 | 0.407 |
| Lymphoma | 0.795 | 0.932 | 0.786 | 0.889 | 0.863 | 0.915 | 0.573 |
| Glomus tumour | 1.000 | 1.000 | 1.000 | 1.000 | 1.000 | 1.000 | 1.000 |
| Intramuscular myxoma | 0.576 | 0.273 | 0.273 | 0.788 | 0.394 | 0.606 | 0.424 |
| Ewing sarcoma | 0.615 | 1.000 | 0.846 | 0.692 | 1.000 | 0.923 | 0.385 |
| Schwannoma | 0.688 | 0.922 | 0.922 | 0.896 | 0.909 | 0.883 | 0.494 |
| Myxoid liposarcoma | 0.795 | 0.923 | 0.897 | 0.974 | 0.974 | 0.923 | 0.256 |
| Leiomyosarcoma | 0.875 | 0.922 | 0.859 | 0.938 | 0.938 | 0.938 | 0.672 |
| Solitary fibrous tumour | 0.607 | 0.832 | 0.842 | 0.816 | 0.842 | 0.633 | 0.709 |
| Low grade fibromyxoid sarcoma | 0.889 | 0.921 | 0.794 | 0.873 | 0.889 | 0.667 | 0.825 |
| **Overall accuracy** | **0.721** | **0.847** | **0.772** | **0.860** | **0.852** | **0.782** | **0.533** |

| Trained with internal and external sets (Cohort 1-Train and Cohort 4-Train), tested on internal set (Cohort 1-Test) |  |  |  |  |  |  |  |
| --- | --- | --- | --- | --- | --- | --- | --- |
| Subtype | Google | UNI-v2 | CONCH-v1.5 | TITAN | Virchow | PRISM | ResNet50 |
| Dermatofibrosarcoma protuberans | 0.850 | 0.850 | 0.900 | 0.950 | 0.950 | 0.850 | 0.750 |
| Neurofibroma | 0.700 | 1.000 | 0.900 | 0.700 | 1.000 | 0.800 | 0.700 |
| Nodular fasciitis | 0.900 | 0.900 | 0.800 | 0.950 | 0.944 | 0.850 | 0.800 |
| Desmoid Fibromatosis | 0.900 | 0.800 | 0.850 | 0.900 | 0.900 | 0.900 | 0.800 |
| Synovial sarcoma | 0.950 | 0.950 | 0.800 | 0.900 | 0.950 | 0.950 | 0.900 |
| Lymphoma | 0.900 | 1.000 | 1.000 | 1.000 | 1.000 | 0.950 | 0.900 |
| Glomus tumour | 1.000 | 1.000 | 1.000 | 1.000 | 1.000 | 1.000 | 0.900 |
| Intramuscular myxoma | 1.000 | 1.000 | 1.000 | 1.000 | 0.950 | 1.000 | 0.900 |
| Ewing sarcoma | 0.950 | 0.950 | 0.950 | 1.000 | 1.000 | 1.000 | 0.800 |
| Schwannoma | 0.900 | 0.900 | 0.850 | 0.900 | 0.900 | 0.900 | 0.800 |
| Myxoid liposarcoma | 0.850 | 0.900 | 0.800 | 1.000 | 0.850 | 0.800 | 0.800 |
| Leiomyosarcoma | 0.800 | 0.800 | 0.850 | 0.850 | 0.850 | 0.750 | 0.900 |
| Solitary fibrous tumour | 0.900 | 0.900 | 0.950 | 0.950 | 0.900 | 0.800 | 0.850 |
| Low grade fibromyxoid sarcoma | 1.000 | 0.900 | 1.000 | 0.950 | 1.000 | 0.850 | 0.950 |
| **Overall** | **0.900** | **0.918** | **0.904** | **0.932** | **0.942** | **0.886** | **0.839** |

| Trained with internal and external sets (Cohort 1-Train and Cohort 4-Train), tested on external set (Cohort 4-Test) |  |  |  |  |  |  |  |
| --- | --- | --- | --- | --- | --- | --- | --- |
| Subtype | Google | UNI-v2 | CONCH-v1.5 | TITAN | Virchow | PRISM | ResNet50 |
| Dermatofibrosarcoma protuberans | 0.946 | 0.973 | 0.811 | 0.973 | 0.919 | 0.838 | 0.757 |
| Neurofibroma | 0.872 | 0.974 | 0.897 | 0.897 | 0.923 | 0.795 | 0.667 |
| Nodular fasciitis | 0.706 | 0.882 | 0.529 | 0.882 | 0.765 | 0.765 | 0.235 |
| Desmoid Fibromatosis | 0.824 | 0.882 | 0.824 | 0.912 | 0.882 | 0.912 | 0.618 |
| Synovial sarcoma | 0.654 | 0.790 | 0.654 | 0.790 | 0.753 | 0.778 | 0.630 |
| Lymphoma | 0.897 | 0.923 | 0.915 | 0.932 | 0.983 | 0.906 | 0.923 |
| Glomus tumour | 1.000 | 1.000 | 1.000 | 1.000 | 1.000 | 1.000 | 1.000 |
| Intramuscular myxoma | 0.970 | 0.515 | 0.788 | 0.879 | 0.697 | 0.788 | 0.697 |
| Ewing sarcoma | 0.769 | 1.000 | 0.923 | 0.846 | 0.923 | 0.923 | 0.538 |
| Schwannoma | 0.792 | 0.935 | 0.909 | 0.883 | 0.779 | 0.870 | 0.740 |
| Myxoid liposarcoma | 0.872 | 0.974 | 0.949 | 1.000 | 0.923 | 0.923 | 0.846 |
| Leiomyosarcoma | 0.891 | 0.969 | 0.906 | 0.938 | 0.922 | 0.938 | 0.797 |
| Solitary fibrous tumour | 0.704 | 0.918 | 0.842 | 0.827 | 0.842 | 0.658 | 0.709 |
| Low grade fibromyxoid sarcoma | 0.873 | 0.873 | 0.762 | 0.857 | 0.889 | 0.714 | 0.746 |
| **Overall** | **0.808** | **0.899** | **0.838** | **0.881** | **0.865** | **0.803** | **0.736** |

## Appendix C Extended figures

**Fig. C3.**
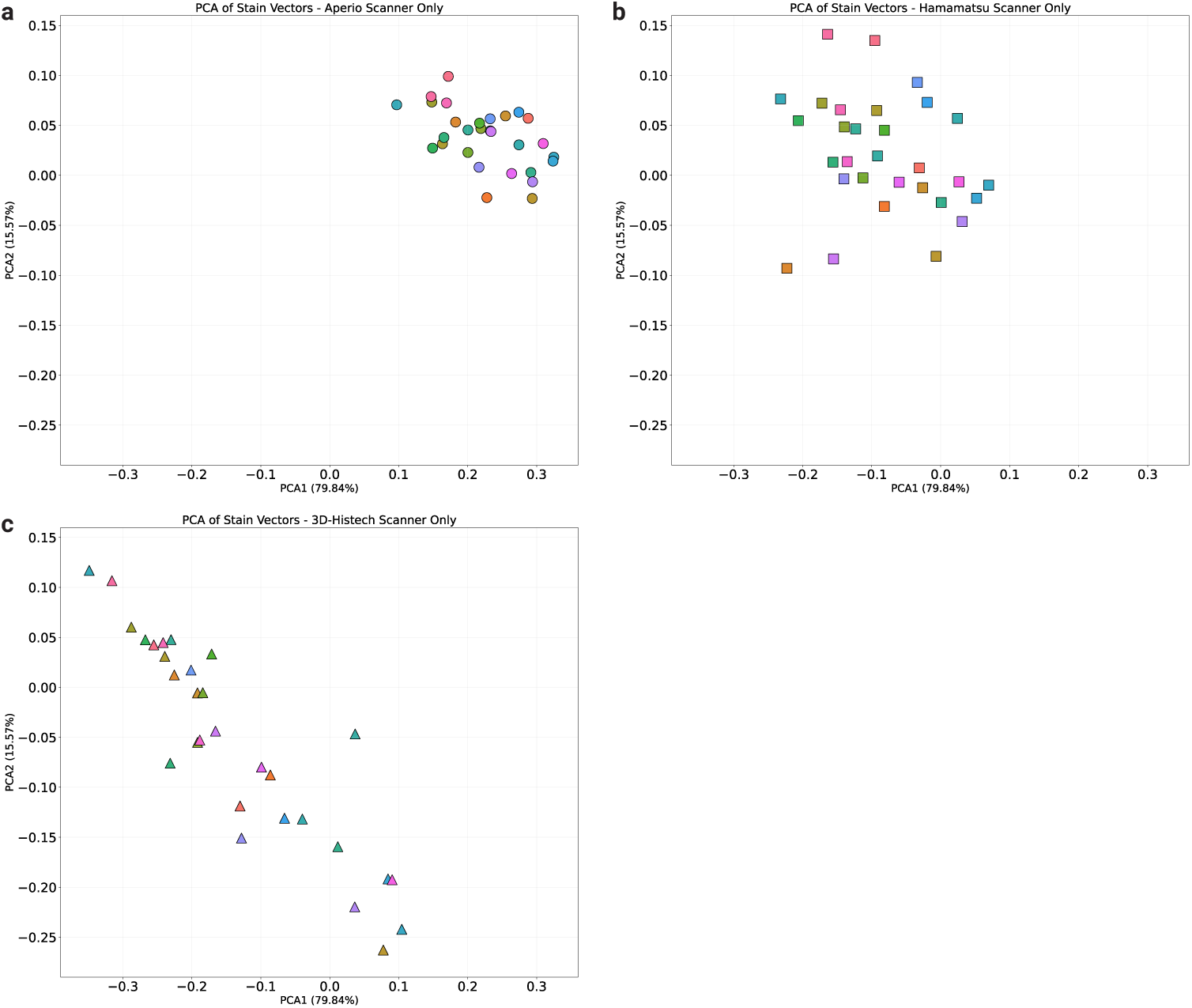
Principal component analysis (PCA) of stain vectors extracted from each 3000 *×* 3000 pixel tile, showing the plot by each scanner: Aperio (**a**), Hamamatsu (**b**), and 3D-Histech (**c**).

